# ANME-2a drive methane oxidation in brackish coastal sediments via multiple pathways

**DOI:** 10.64898/2026.05.06.723182

**Authors:** Robin Klomp, Anna J. Wallenius, Merijn A.W. Schutgens, Theo van Alen, Thomas Röckmann, Mike S.M. Jetten, Caroline P. Slomp

**Affiliations:** Department of Microbiology, Radboud Institute for Biological and Environmental Sciences, Radboud University, Heyendaalseweg 135, 6525AJ Nijmegen, the Netherlands; Department of Earth Sciences, Utrecht University, Princetonlaan 8a, 3584 CB Utrecht, the Netherlands; Institute for Marine and Atmospheric Research Utrecht, Utrecht University, Princetonplein 5, 3584CC Utrecht, the Netherlands

## Abstract

Methane is a powerful greenhouse gas. Typically, a large fraction of the methane formed in coastal sediments is removed via anaerobic methane oxidation (AOM). Here, we demonstrate the potential for a range of AOM pathways in brackish coastal sediments by ANME-2a archaea. At our study site, geochemical profiles indicate that AOM is primarily restricted to a shallow, metal-oxide-rich sulfate-methane transition zone (SMTZ). ANME-2a were the sole methanotrophs detected, and metatranscriptomics showed the highest expression levels of the ANME-2a genes in the SMTZ. AOM activity was observed in sediment incubations with various electron acceptors, including sulfate, metal oxides, and the organic matter analogue graphene oxide. Highest potential rates were observed in sediments from below the SMTZ, pointing towards fast stimulation of the deeper methanotrophic community when alleviating the electron acceptor limitation. The variety of AOM pathways and persistence of methanotrophs below the SMTZ likely contribute to the resilience of the microbial methane filter in brackish coastal sediments.

## 1. Introduction

Methane is a key greenhouse gas that, in marine sediments, is mainly produced by methanogenic archaea (Reeburgh, 2007). In most continental margin sediments, the methane is efficiently removed through anaerobic and aerobic oxidation (Knittel & Boetius, 2009). In coastal sediments, however, some of the methane may escape oxidation, especially when the zone of methanogenesis approaches the sediment-water interface (Lapham et al., 2024; Żygadłowska et al., 2024). As a consequence, coastal regions may contribute up to 75% of total marine methane emissions, even though these systems account for only a small part of the oceanic surface area (Hamdan & Wickland, 2016). This makes it crucial to understand the microbial pathways contributing to methane removal in coastal sediments.

Coastal sediments typically receive high inputs of organic matter, which induce sharp redox gradients near the sediment-water interface (Burdige, 2007). Besides oxygen, which generally penetrates down to depths of only a few millimeters, other electron acceptors used to degrade organic matter are, in order of decreasing energy yield, nitrate/nitrite, manganese (Mn) oxide, iron (Fe) oxide and sulfate (SO_4_^2-^; Froelich et al., 1979). When these electron acceptors are depleted, fermentation and methanogenesis are the remaining pathways (Canfield & Thamdrup, 2009). Most of the methane is thought to be removed through anaerobic oxidation of methane (AOM) with sulfate as the electron acceptor (S-AOM) in the so-called sulfate-methane transition zone (SMTZ) (Boetius et al., 2000; Orphan et al., 2002; Reeburgh, 2007).

In environments subject to non-steady state deposition, methane may also come into contact with metal oxides, creating an opportunity for a coupling between manganese and iron oxide reduction and AOM (Mn- and Fe-AOM, respectively; Wallenius et al., 2021 and references therein). The first evidence for potential Mn- and Fe-AOM was obtained from incubations of marine methane-seep sediments using ^13^C-labelled methane (Beal et al., 2009). Follow-up experimental and modeling studies confirmed a potential role for Mn- and/or Fe-AOM in brackish and marine sediments (Segarra et al., 2013; Riedinger et al., 2014; Egger et al., 2015; Aromokeye et al., 2020; Lenstra et al., 2023, Klomp et al., 2026). Brackish sediments where an increased input of organic matter has led to an upward shift of the SMTZ are particularly conducive to Mn- and Fe-AOM, because of the presence of metal oxides in a zone with high methane concentrations (Egger et al., 2015; Rooze et al., 2016; Lenstra et al., 2023). Besides direct coupling of metal oxide reduction to AOM in such settings, metal oxides may also drive S-AOM via the production of sulfate in a cryptic sulfur cycle (Holmkvist et al., 2011; Norði et al., 2013; Su et al., 2020, Klomp et al., 2026). Since there is no conclusive geochemical signature for Mn-AOM, Fe-AOM or cryptic sulfur cycling, the need to identify the microorganisms involved is important.

Typically, S-AOM is performed by a consortium of anaerobic methanotrophic archaea (ANME) and sulfate-reducing bacteria (SRB) (Boetius et al., 2000; Orphan et al., 2002). The clade *Ca.* Methanoperedenaceae (also known as ANME-2d) has been linked to Mn- and Fe-AOM reduction in freshwater sediments (Ettwig et al., 2016; Cai et al., 2018; Leu et al., 2020). In brackish and marine sediments, ANME-2d is typically only present at very low abundances and thus is unlikely to contribute to Mn- and Fe-AOM (Aromokeye et al., 2020; Rasigraf et al., 2020; Wallenius et al., 2021, 2025). Other subclades of ANME-2 may be involved instead, such as ANME-2ab, now classified as *Ca.* Methanocomedenaceae (Chadwick et al., 2022). ANME-2a was, for example, proposed to be linked to Fe-AOM based on sediment incubations for a marine site in the North Sea (Aromokeye et al., 2020). Similar depth trends in ANME-2a and total Fe contents (Rasigraf et al., 2020) for sediments from the brackish Bothnian Sea, for which Fe-AOM activity was reported earlier (Egger et al., 2015), also point to a potential role for ANME-2a in Fe-AOM. Finally, AOM may also be coupled to redox-active humic substances in natural organic matter (NOM-AOM) (Scheller et al., 2016; Valenzuela et al., 2017; Zhao et al., 2024). While in freshwater sediments, ANME-2d can be involved in NOM-AOM (Bai et al., 2019), various other ANME clades have been inferred to carry out the process in brackish and marine environments, including ANME-1b (anoxic marine water; van Grinsven et al., 2020), ANME-2a and 2c (marine methane seep; Scheller et al., 2016) and ANME-2ab (brackish canal sediment; Pelsma et al., 2023).

In this study, we examine the potential and *in-situ* relevance of various AOM pathways and the microbes involved in methane- and metal oxide-rich sediments from a brackish coastal sea (Bothnian Sea). We focus on sediments from within and below the SMTZ and assess the following electron acceptors in AOM: sulfate, Mn and Fe oxides and natural organic matter. A complementary set of methods was used, including geochemical profiling of sediment and pore water, 16S rRNA gene amplicon sequencing, metagenomics, metatranscriptomics, and sediment incubations using ^13^C-labelled methane. We find evidence for a range of AOM pathways, performed by ANME2-a, with the highest *in-situ* activity in the SMTZ but the greatest potential activity deeper in the sediment, indicative of a resilient and easily reinvigorated methanotrophic community.

## 2. Materials and methods

### 2.1 Study site

The Bothnian Sea is a brackish oligotrophic coastal basin in the northern part of the Baltic Sea. Rivers are the main source of organic matter in the region. Inputs of this terrestrial organic matter vary greatly between years, linked to variations in rainfall and river discharge, and appear to be increasing, linked to climate change (Algesten et al., 2006; Lenstra et al., 2018). Land uplift in the Bothnian Sea region contributes to the transport of this material from shallow areas to the deeper basins (Leivuori & Niemistö, 1995). The input of marine organic matter has also gained importance over the past decades as a result of anthropogenic eutrophication (Kuliński et al., 2022). The increase in deposition of both terrestrial and marine organic matter is thought to be responsible for the observed shallowing of the SMTZ at many locations in the Bothnian Sea (Slomp et al., 2013; Egger et al., 2015; Rooze et al., 2016). Bothnian Sea sediments are metal oxide-rich, with frequent burial of metal oxides into and below the SMTZ (Slomp et al., 2013; Egger et al., 2015; Lenstra et al., 2018).

The sampling location, US2 (62°50.99’ N; 18°53.53’ E; Fig. S1), has a permanently oxygenated water column, a sediment accumulation rate of 1.6 cm yr^-1^ and is characterized by a high input of metal oxides and a shallow SMTZ (Slomp et al., 2013). Macrofaunal activity, mainly by burrowing crustaceans and polychaetes, affects the sediment via bioturbation and bioirrigation (Josefson et al., 2012).

### 2.2 Sediment and pore water collection

In May 2022, six sediment cores were collected on board Research Vessel KBV 181, using a Gemini corer system with transparent core liners of 8 cm inner diameter: one core for porewater and solid phase analysis, one for methane, one for sulfate reduction rates, one for porosity and DNA/RNA samples and two to collect material for incubation experiments.

For pore water and solid phase analysis, two bottom water samples were taken. The core was sectioned under a N_2_ atmosphere in a glove bag at a 1 cm depth resolution until 10 cm depth and a 2 cm depth resolution below 10 cm depth. The sediment was transferred into 50 mL centrifuge tubes and centrifuged at 4500 rpm for 20 minutes to separate the sediment from the pore water. The sediment was stored in N_2_ purged aluminum bags at -20°C until further processing for solid phase analysis. The supernatant was filtered (0.45 µm) in a N_2_ filled glove bag and subsampled for analysis of alkalinity, sulfate, ammonium (NH_4_^+^), dissolved Mn, Fe, phosphorus (P), sulfide (H_2_S) and dissolved inorganic carbon (DIC), including the DIC C-isotopic signature. Aliquots for alkalinity and sulfate were stored in polyethylene vials at 4°C until analysis. Samples for ammonium analysis were stored at -20°C. Samples for total dissolved Mn, Fe and P were acidified with 10 µL 30% suprapur HCl per milliliter of sample and stored at 4°C. Samples for sulfide analysis were diluted 5 times in 2% Zn acetate in a glass vial and stored at 4°C. Samples for DIC and DIC C-isotopic signal were stored in airtight glass vials without headspace and with 10 µL HgCl_2_ per 2 ml sample at 4°C.

Samples for the determination of methane, including C- and D-isotopes, and sulfate reduction rates were taken directly upon recovery from a core liner with pre-drilled holes at a 2.5 cm depth resolution covered with tape prior to coring. For methane, 10 ml of sediment was transferred directly into a 65 ml glass bottle filled with a saturated NaCl solution using pre-cut plastic syringes. Bottles were immediately stoppered, capped and stored upside down until analysis. For sulfate reduction rates, 5 ml of sediment was sampled using pre-cut syringes, which were sealed off with parafilm and stored in a N_2_ flushed aluminum bag at 4°C.

For porosity and DNA/RNA sample collection, a core was sectioned under atmospheric conditions in sections of 1 cm in the top 10 cm of the core and in sections of 2 cm below 10 cm. Part of the sediment was placed in pre-weighed centrifuge tubes and stored at 4°C until porosity analysis. Another part was conveyed to autoclaved Eppendorf tubes, frozen with liquid N_2_ and stored at -20°C for DNA and RNA analyses.

The remaining cores were stored at 4°C and sectioned within two weeks after retrieval under an N_2_ atmosphere at a 4 cm resolution. The sediment was placed in plastic bags and stored in N_2_ flushed aluminum bags at 4°C until use in sediment incubations.

### 2.3 Chemical analysis of solid phase

Sample residues for the solid phase analysis were freeze-dried and ground with an agate mortar and pestle, all under a N_2_ atmosphere. Aliquots of the sediment (∼300 mg) were used to determine C_org_ contents. The sediment was decalcified with 1 M HCl (via a two-step wash; Van Santvoort et al., 2002), dried, weighed and powdered in an agate mortar and pestle. The powdered sediment was analyzed using an elemental analyzer (Fison Instruments model NA 1500 NCS). The C content was corrected for weight loss during decalcification and an internationally certified soil standard IVA2 was used to determine the accuracy and precision of the analysis. The certified C content in IVA2 is 0.732 wt%, the measured mean value for C in IVA2 (n = 7) was 0.716 wt% with a standard deviation of 0.004 wt%.

Another aliquot (∼50 – 100 mg) was used to determine the mineral phases of Mn and Fe via a sequential extraction. For Mn, we used the extraction scheme of Lenstra et al. (2021), for Fe, the procedure described by Kraal et al. (2017). Details are provided in supplemental Table S1 and S2. In the first step of the Fe extraction (1 M HCl), Fe(II) and Fe(III) were measured separately via the spectrophotometric method using 1-10 phenantroline (APHA, 2005), to distinguish Fe-oxides from reduced Fe minerals such as FeS and Fe carbonate. In the other steps, extracted Mn and Fe was measured via inductively coupled plasma – optical mission spectroscopy (ICP-OES; PerkinElmer Avio; detection limit 0.1 µmol L^−1^ for Fe and 0.02 µmol L^−1^ for Mn). The standard deviation over all extraction steps based on duplicates (n = 3) was 2.5% for Fe and 2.4% for Mn.

Iron sulfide (FeS) concentrations were determined via the passive diffusion method as described by (Burton et al., 2008) on sediment aliquots of ∼300 mg. The standard deviation based on duplicates (n = 3) was 5.0%.

Total sediment Mn, Fe, P and S was determined via a sediment digestion of ∼100 mg of sediment in 2.5 ml mixed acid (HNO_3_; HClO^-^; 2:3) and 2.5 ml 40% HF at 90°C. After evaporation of the acid mixture, the residue was redissolved in 1 M HNO_3_. The solution was analyzed using ICP-OES with the following recoveries: 106% for Mn, 107% for Fe, 104% for P and 101% for S. The standard deviation (n=2) was 1% for Mn, 1.1% for Fe, 1.1% for P and 2.1% for S. Porosity was determined based on the weight loss of the sample after drying in an oven at 60°C.

### 2.4 Chemical analysis of pore water

Porewater Fe, Mn and P were measured via ICP-OES (PerkinElmer Avio; detection limit 0.1 µmol L^−1^ for Fe, 0.02 µmol L^−1^ for Mn, 3.1 µmol L^−1^ for P). A 10 mL N_2_ headspace was injected into the methane bottles and after an equilibration time of 7 days, the headspace was analyzed on a Thermo Finnigan Trace™ gas chromatograph (flame ionization detector; limit of detection 0.02 µmol L^−1^). The isotopic composition of methane (δ^13^C-CH_4_ and δD-CH_4_) was analyzed by Continuous Flow Isotope Ratio Mass Spectrometry (CF-IRMS) as described in Brass & Röckmann (2010) and Sapart et al. (2011). Porewater sulfate was determined via ion chromatography (IC; Metrohm 930 Compact IC Flex; detection limit of 10 µmol L^−1^). Porewater sulfide was measured spectrophotometrically using the phenylendiamine and ferric chloride method (Cline, 1969; detection limit of 1 µmol L^-1^). Alkalinity was measured within 24 h after sample collection through titration with 0.01 M HCl. Porewater DIC including its isotopic composition was determined after acidification of the sample with 85% H_3_PO_4_ in an argon purged exetainer and measurement of the headspace on a GasBench – Isotope Ratio Mass Spectrometer (GasBench II Delta V Advantage IRMA, Thermo Scientific). Concentrations of ammonium were determined via the indophenol blue spectrophotometric method (Solórzano, 1968).

### 2.5 Determination of sulfate reduction rates

To determine sulfate reduction rates, the samples were injected with 100 kBq ^35^S-SO_4_^2-^ six days after core retrieval, sealed again and incubated for 24 h in N_2_-purged aluminum bags at 4°C. The sediment was then transferred into a 50 mL centrifuge tube containing 20 mL oxygen-free 20% zinc acetate to precipitate any sulfide formed and terminate microbial activity (Fossing & Jørgensen, 1989; Kallmeyer et al., 2004). The samples were stored at -20°C in aluminum nitrogen purged bags. Upon analysis, the samples were washed twice with oxygen-free bottom water and centrifuged to remove pore water and unreacted ^35^S-SO_4_^2-^ (Egger et al., 2016). The reduced S was then extracted with an acidic chrome chloride solution for 48 h (Kallmeyer et al., 2004) and captured in 20% zinc acetate via the passive diffusion method as described in Burton et al. (2008). The formed radioactive sulfide was determined by mixing the 20% zinc acetate 1:2 vol:vol with Ecoscint XR (NAT1396, Scientific Laboratory Supplies, UK) and analysis on an automatic triple-to-double coincidence ratio (TDCR) liquid scintillation counter (Hidex 600 SL, LabLogic Systems Limited, UK). Sulfate reduction rates were quantified by comparing the activity (decays per minute) of the radiolabeled total reduced inorganic sulfur (α_TRIS_) to the total sulfate radiotracer (α_TOT_) as described in Kallmeyer et al. (2004):

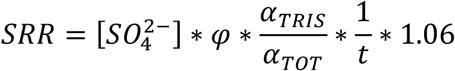

Where 𝜑 is the measured porosity, t is the incubation time in days and 1.06 is the correction factor for the expected isotopic fractionation (Jørgensen & Fenchel, 1974; Kallmeyer et al., 2004).

### 2.6 DNA extraction and 16S rRNA amplicon sequencing and analysis

DNA was isolated with the DNeasy PowerSoil Pro DNA isolation kit (Qiagen, Venlo, Netherlands) according to the manufacturer’s instructions after bead-beating the samples on a TissueLyser LT (Qiagen) for 10 min at 50Hz. Samples from the incubations were pre-incubated at 65°C for 10 min in the kit’s C1 solution before bead-beating to increase archaeal DNA yields. 16S rRNA gene amplicon sequencing was performed on the Illumina MiSeq Next Generation Sequencing platform by Macrogen (Seoul, South Korea) using Herculase II Fusion DNA Polymerase Nextera XT Index Kit V2, yielding 2x300bp paired end reads. Archaeal primers used were Arch349F (5′-GYGCASCAGKCGMGAAW-3′) and Arch806R (5′-GGACTACVSGGGTATCTAAT-3′; Takai & Horikoshi, 2000) and for bacteria Bac341F (5′-CCTACGGGNGGCWGCAG-3′; Herlemann et al., 2011) and Bac806R (5′-GGACTACHVGGGTWTCTAAT-3′; Caporaso et al., 2012). The optimal trimming parameters were determined with Figaro (Weinstein et al., 2019), and trimming and further processing were done using the DADA2 pipeline (v1.8; Callahan et al., 2016) in Rstudio (v4.1.1; R Core Team, 2019) as explained previously (Wallenius et al., 2025). Raw reads are accessible on the National Center for Biotechnology Information (NCBI) website under the accession number PRJNA1306556.

### 2.7 Metagenomic sequencing and analysis

The DNA from sediment depths of 16-20 cm and 28-32 cm was used for short-read metagenomic sequencing using a TruSeq DNA PCR-free library with an insert size of 550bp on NovaSeq6000 (Illumina) platform by Macrogen (Seoul, South Korea), producing 2 × 150bp paired-end reads (100 Gbp/sample). For long read sequencing with Nanopore, library preparation was done starting with 150 to 400 ng of DNA. The quality of the DNA was checked by agarose gel electrophoresis. The DNA Library construction was performed using the Native Barcoding Kit 24 V14 (SQK-NBD114-24), according to the manufacturer’s protocol (Oxford Nanopore Technologies, Oxford, UK). The library was loaded on a Flow Cell (R10.4.1) and sequenced using the MinION Mk1C device (Oxford Nanopore Technologies, Oxford, UK), according to the manufacturer’s instructions. Quality of the sequence reads was analyzed using fastqc (Andrews, 2010). We used singleM (Woodcroft et al., 2024) to characterize the samples taxonomically. The quality control, trimming, assembly and binning was done using the Aviary pipeline (v.0.11.0; Newell et al., 2025). The reads were trimmed with fastp (v0.24.0; Chen et al., 2018), assembled with metaSPAdes (v.4.0.0; Nurk et al., 2017), mapped with CoverM (v0.7.0; Aroney et al., 2025), and binned with MetaBAT andi MetaBAT2 (v2.15; Kang et al., 2015, 2019), Semibin (Pan et al., 2022), Rosella (v0.5.5; https://rhysnewell.github.io/rosella/) and DASTool (v.1.1.2.; Sieber et al., 2018). The metagenomically assembled genomes (MAGs) were taxonomically classified with GTDB-Tk release 220 (v2.4.0; Chaumeil et al., 2022) and the MAG completeness and contamination was estimated with CheckM2 (v1.0.2; Aroney et al., 2025). We also used differential coverage binning (Albertsen et al., 2013) as described previously (Wallenius et al. 2025, 2026) to recover one ANME MAG. Genomes were annotated with DRAM v1.0 (Shaffer et al., 2020) and metascan (Cremers et al., 2022). Genes involved in metal reduction were retrieved with FeGenie (Garber et al., 2020). Proteins were identified with prodigal (v2.6.3; Hyatt et al., 2010) and used to search for proteins with heme-binding motifs (>3).

### 2.8 Metatranscriptomics

RNA was extracted from freeze-dried sediment from depths of 16-20 cm and 28-32 cm with the RNeasy PowerSoil Total RNA Kit (Qiagen, Hilden, Germany), using the protocol with the following modifications to increase the RNA yield: the input amount was increased to ∼0.5 g sediment, in step 2 we added an extra 0.5 ml DEPC-treated water (Invitrogen, Carlsbad, United States) to increase the volume of the aquatic phase; in step 3 we added 4 ml of phenol/chloroform/isoamyl alcohol to counter-act for the increased humic acids; in step 9, one ml DEPC treated water was added before incubating 30 min at -20°C, and the final extraction was done with 25 µl SR7 volume to increase the RNA concentration. DNA contamination was removed with Invitrogen DNA removal kit (Fischer Scientific, USA). The quality of the RNA was checked using the 2100 Bioanalyzer instrument (Agilent technologies, USA) and the Agilent RNA 6000 Nano Kit according to the manufacturer’s instructions. Metatranscriptomic sequencing was performed using a TruSeq stranded with NEB rRNA depletion kit (bacteria) (Illumina, USA) on a NovaSeqX (Illumina) platform by Macrogen (Seoul, South Korea), generating 150-bp paired-end reads with ∼15 Gb throughput/sample. For both depths three biological replicates per depth from different extractions were sequenced. Raw sequences were quality trimmed and filtered for rRNA encoding reads, mapped against the DRAM-generated scaffolds, and transcripts permillion (TPM) values were generated using transcriptm v0.5 (https://github.com/sternp/transcriptm). Relative abundance was calculated using CoverM v0.6.1. Further analysis, including data visualization was done in RStudio v4.4.0. Map reads corresponding to coding DNA sequences (CDS) were converted into transcripts per million and pooled per genome to compare the relative abundance of MAGs expressed per sample.

### 2.9 Incubation experiment

The methane oxidation potential of various electron acceptors, namely oxygen, sulfate, Mn oxide, Fe oxide and graphene oxide, as an analogue for humic acid, was tested via batch incubations using sediment from within the SMTZ (16 - 20 cm) and below the SMTZ (28 - 32 cm). For both intervals, methane and metal oxides overlapped in-situ. The incubations were started within six months of sediment collection.

The sediment was diluted in a 1:4 ratio with HEPES buffered artificial sulfate-free seawater (ASW; for composition see table S3) under an anoxic atmosphere. The sediment slurry was distributed over sterile 120 mL serum bottles, to a final amount of 60 g slurry per bottle. In the incubations with sulfate, 10 mmol L^-1^ of Na_2_SO_4_ solution was added. Manganese was added in the form of birnessite (7.5 mmol L^-1^; MnO_2_\**n*H_2_O) synthesized according to the protocol by Händel et al. (2013) and iron was added in the form of ferrihydrite (7.5 mmol L^-1^; FeO(OH)) synthesized as described in Raven et al. (1998). Both metal oxides were analyzed via X-ray diffraction to characterize the minerals (Fig. S2). Graphene oxide (C_x_H_y_O_z_) was added in a concentration of 200 mg L^-1^, after diluting graphene oxide paste (Merck KGaA, Darmstadt, Germany) in demineralized water and deoxygenating the solution. The bottles were closed with capped rubber stoppers and, in the incubation with oxygen, air was injected to achieve 4% oxygen in the headspace. The sediment was first pre-incubated for one week, shaking at 90 rpm at room temperature with 100% N_2_ headspace, to oxidize sediment FeS and minimize the release of sulfate in a cryptic sulfur cycle in the real incubations. After one week, the ASW was replaced by freshly prepared ASW under an anoxic atmosphere and new electron acceptors were added to each bottle, in the same concentrations as in the pre-incubations. The headspace was replaced by a headspace of 76% N_2_, 4% CO_2_ and 20% ^13^C-labelled methane (Cambridge Isotope Laboratories, Inc., Andover, USA). The headspace in the O_2_-MO incubation was amended with 50% air, 4% CO_2_ and 20% ^13^C-CH_4_. The bottles were then placed at room temperature shaking at 90 rpm for 94 days. In the O_2_-MO incubation, new oxygen was added in the form of pure oxygen gas to a concentration of 4% of the headspace after 28 and 49 days. In the NOM-AOM incubations with sediment from below the SMTZ, new substrate was added to a concentration of 200 mg L^-1^ after 56 days, when the activity stagnated likely due to substrate depletion.

The headspace was measured weekly for ^12^C-CO_2_ and ^13^C-CO_2_ concentrations on a gas-chromatograph coupled to a mass spectrometer (GC-MS; Agilent 5975C inert MSD, detection limit for accurate CO_2_ isotope concentrations is 40 nmol CO_2_) to monitor ^13^C-CH_4_ oxidation. Upon addition of Mn oxides, the headspace concentration of CO_2_ decreased, possibly because CO_2_ was drawn into the dissolved phase upon an increase in the pH during Mn oxide reduction (Silburn et al., 2017). To prevent CO_2_ concentrations below the detection limit, new CO_2_ (4% of the headspace) was added to the Mn-AOM incubation with sediment from below the SMTZ after 49 days.

Every other week, a sample was taken from the medium with a N_2_ purged syringe and needle through the stopper, which was used for the analysis of dissolved Mn and Fe (via ICP-OES, iCAP 6300 with detection limits of 0.02 and 0.18 µmol L^-1^ for Mn and Fe, respectively), sulfate (via IC), sulfide and dissolved Mn(II) and dissolved Mn(III) (spectrophotometrically, as described in Madison et al. (2011); Oldham et al. (2015)). Every four weeks, a sample was taken to determine dissolved CO_2_, including the CO_2_ isotopes. A liquid sample of 0.5 ml was injected in an argon flushed 3 ml exetainer containing 1 ml 85% phosphoric acid and the headspace of the exetainer was injected into the GC-MS.

Samples for 16S rRNA amplicon sequencing were obtained at the start of the pre-incubations and from each bottle separately at the end of the incubation experiment of sediment from below the SMTZ (28 - 32 cm). These samples were processed as described in section 2.6.

## 3. Results

### 3.1 Methane oxidation in a shallow SMTZ

A distinct SMTZ was identified between 12 and 22 cm depth in the sediment at our study site (US2), characterized by strong counter gradients of sulfate and methane in the porewater (Fig. 1). A small fraction of the methane passed onwards to the zone above the SMTZ. Methane isotopic values of δ^13^C and δD were stable below the SMTZ, at values around -79‰ and - 265‰, respectively, and increased in the SMTZ to values of -61‰ and -93‰ (Fig. 1), respectively, indicating methane oxidation. Above the SMTZ, methane isotopic values were stable, apart from the uppermost 2 cm where both values increased.

**Fig. 1.**
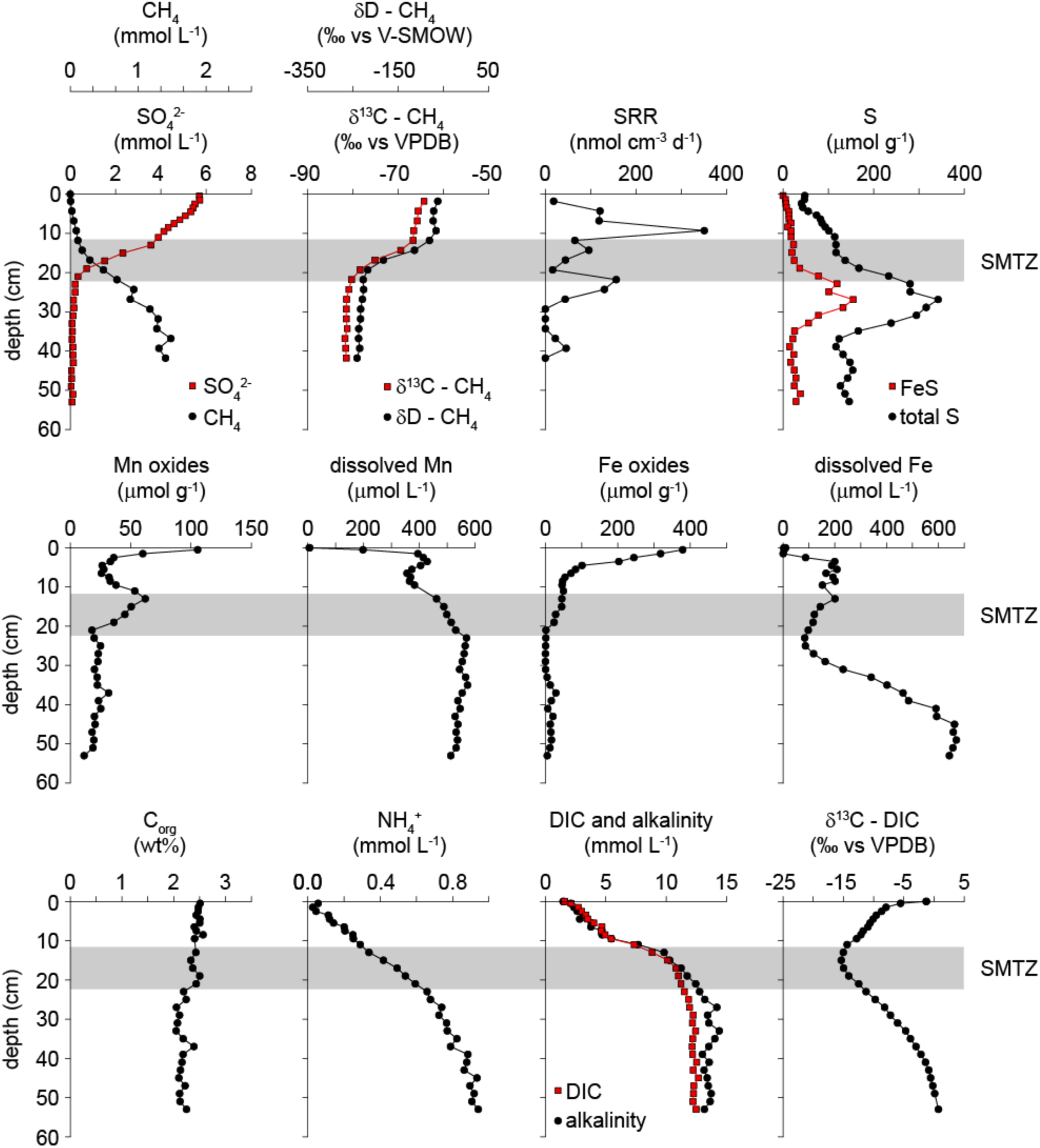
Pore water and sediment depth profiles and sulfate reduction rates (SRR) for our study site US2. The zone indicated with gray shading indicates the SMTZ. The step used to determine Mn oxides may include Mn phosphates. Additional sediment Mn and Fe forms are shown in supplemental Fig. S4.

Sulfate reduction rates were highest just above the SMTZ at a depth of 9 cm, reaching values of up to 352 nmol cm^-3^ d^-1^ (Fig. 1). A second maximum in sulfate reduction rates was observed at the bottom of the SMTZ at a depth of 22 cm, reaching values of up to 157 nmol cm^-3^ d^-1^. Free sulfide was always below the detection limit in the porewater. Below the SMTZ, solid phase total S reached a maximum concentration of 342 µmol g^-1^ at 27 cm depth. A substantial part of the peak in total S, up to 155 µmol g^-1^, consisted of FeS (Fig. 1).

Surface sediments were enriched in Mn and Fe oxides, reaching concentrations of up to 106 and 379 µmol g^-1^, respectively (Fig. 1). At the upper boundary of the SMTZ, a peak in Mn oxides was present, which may partly consist of Mn phosphates dissolved in the same extraction step as the Mn oxides (Fig. S3 and table S1). Within the SMTZ, Mn oxide contents decreased to around 20 µmol g^-1^ and Fe oxide contents decreased from 46 µmol g^-1^ to 0 µmol g^-1^. Mn oxides remained around 20 µmol g^-1^ but Fe oxides emerged again below 29 cm depth, reaching concentrations of up to 29 µmol g^-1^. Dissolved Mn and Fe increased rapidly in the upper few cm of the sediment to around 400 and 200 µmol L^-1^, respectively, and subsequently decreased, but at different depths (Fig. 1). Deeper in the sediment the concentrations of both Mn and Fe increased again, with dissolved Mn and Fe reaching maxima of 566 µmol L^-1^ and 670 µmol L^-1^, respectively.

The C_org_ content was ∼2.5 wt% in the top 20 cm of the sediment and decreased to around 2 wt% below a depth of 20 cm (Fig. 1). Concentrations of ammonium, DIC and alkalinity increased strongly in the top 20 cm of the sediment, indicating the highest rates of organic matter degradation in this zone (Fig. 1). Below 20 cm, ammonium continued to increase with depth, whereas DIC and alkalinity concentrations stabilized around 12 and 13 mmol L^-1^, respectively. Values of δ^13^C of DIC were close to -5.5 ‰ near the sediment water interface and decreased to a minimum of -15.4‰ within the SMTZ (Fig. 1). Below the SMTZ, the δ^13^C of DIC increased to 0.7‰. Porosity decreased from 0.95 at the sediment surface to ∼0.90 at 5 cm depth and remained near this value at greater depth (Fig. S3).

### 3.2 Diversity of the microbial community

The 16S rRNA gene amplicon sequences revealed distinct communities of both archaea and bacteria, which in some cases were related to the boundaries of the SMTZ (Fig. 1 and 2). Ammonia-oxidizing archaea, unclassified *Nitrosopumilaceae* and ‘*Candidatus* Nitrosopumilus’, covered > 65% of all archaeal reads in the top 12 cm. In the SMTZ (12 to 22 cm), ANME-2ab was the only methanotroph detected at a maximum abundance of 31%. No ANME-2b (*Candidatus* Methanomarinus) were found in the metagenomic analysis (Table S4), therefore we will refer to ANME-2a in the following text. Surprisingly, ANME-2a even accounted for up to 61% of all archaeal reads in the interval from 24-34 cm below the SMTZ. We detected four abundant ANME-2a ASVs with slightly different niches, with ASV_1 dominating down to a depth of 34 cm, whereas the other ASVs were either more variable or restricted to the sediments below the SMTZ (Fig. S5). Below 28 cm, methanogenic genera *Methanosarcina*, *Methanoregula* and *Methanosaeta* together with Bathyarchaeaia, appeared and became abundant from a depth of 34 cm onwards, indicating active methane production.

**Fig. 2:**
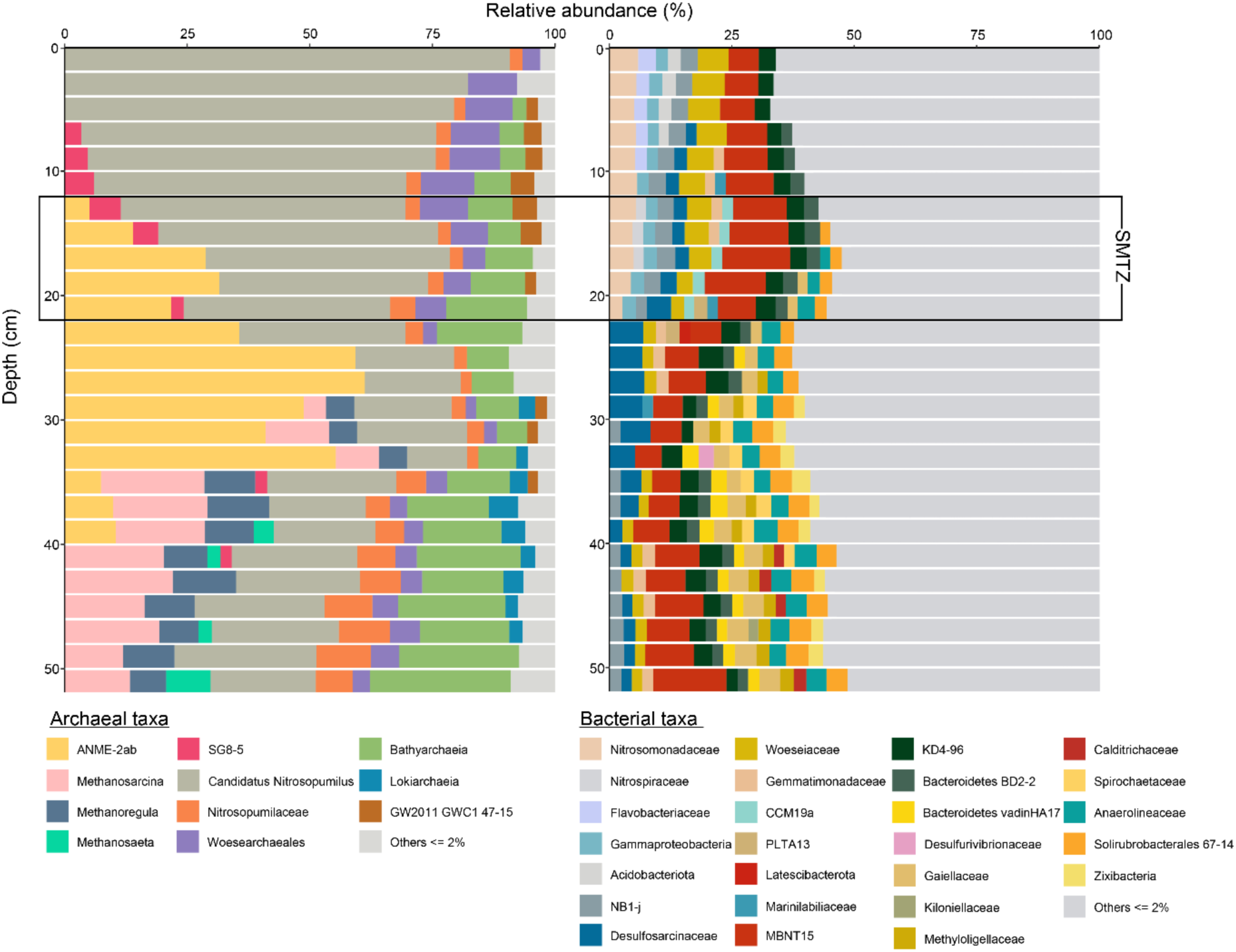
16S rRNA gene amplicon sequencing relative abundances of archaeal DNA (left) and bacterial DNA (right). All taxa with < 2% abundance are grouped in ‘Others’.

The bacterial community was highly diverse, with over 50% of taxa having a relative abundance of less than 2%. Despite this diversity, there was a clear difference in the most abundant taxa with sediment depth. Above and in the SMTZ (0 to 22 cm), *Nitrosomonadaceae*, *Nitrospiraceae, Flavobacteriaceae*, *Woeseiaceae*, unclassified Gammaproteobacteria, and the unclassified phylum NB1-j made up about 25% of all bacterial reads. Between 18 and 34 cm depth, the SRB *Desulfosarcinaceae* was the most abundant bacterial group after the MBNT15 phylum, reaching up to 7% in relative abundance. Below 32 cm, bacterial diversity increased, with *Anaerolineaceae* and S*olirubrobacterales* being the most abundant taxa.

### 3.3 Metabolic potential for sulfur, C1 and metal cycling

We sequenced the metagenomes from two depths, 16-20 cm (SMTZ) and 28-32cm (below the SMTZ), and in total 479 medium-quality (>50% completeness, <10% contamination) MAGs were obtained. These MAGs covered 38% (SMTZ) and 34 % (below the SMTZ) of the total metagenomic reads, thus a significant part of the community was not assembled and thus not analyzed. Based on the mapping of metagenomic reads (Fig. 3), the MAG with the highest relative abundance at both depths was *Defferimicrobium* (MAG US2_17), which accounted for 4.4% of reads in the SMTZ and 2.0% below the SMTZ.

**Fig. 3.**
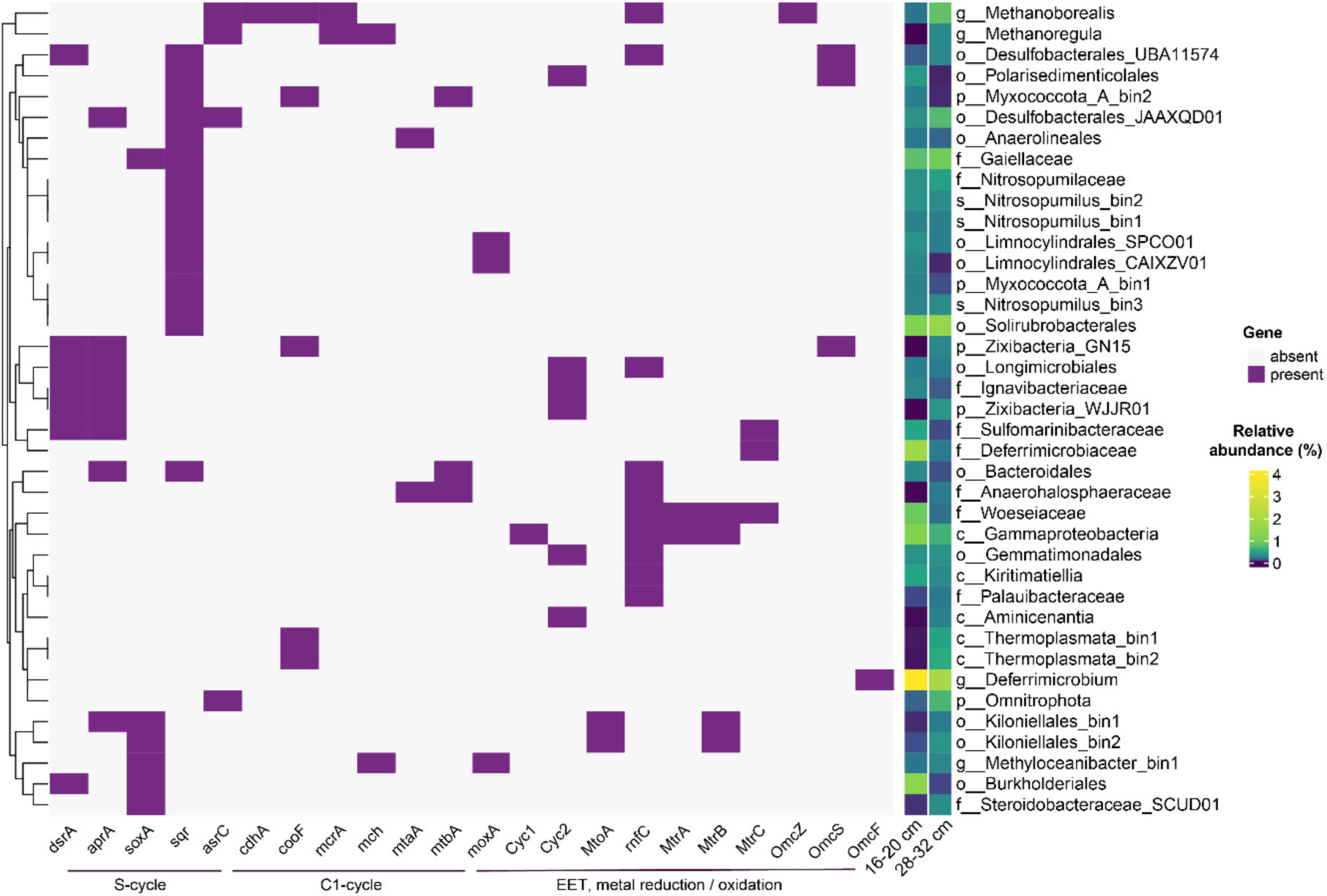
The metabolic potential of the most abundant MAGs (> 0.2 % abundance at one depth) based on S-cycling genes (*dsr*A, *apr*A, *sox*A, *sqr*, *asr*C), C1-cycling (*cdh*A, *coo*F, *mcr*A, *mch*, *mta*A, *mtb*A, *mox*A) and genes involved in metal reduction/oxidation or EET (*cyc*1, *cyc*2, *mto*A, *mtr*A, *mtr*C, *omc*Z, *omc*S, *omc*F). The MAGs are named based on the lowest taxonomic rank with a classified name. The relative abundance of each MAG at the two depths is indicated in the two columns before the genome name.

As seen from the 16S rRNA gene amplicon profile of the sediment, the most abundant taxa differed significantly between the two depths. In the SMTZ, *Deferrimicrobiaceae* (US2_15), Burkholderiales (US2_35), Solirubrobacterales and Gammaprotebacteria (US2_34) were the only bins with >1 % abundance. The sediment below the SMTZ was more diverse and only Deferrimicrobium and Solirubrobacterales reached >1 % abundance. Curiously, MAGs representing likely SRBs (phylum Desulfobacterota) were higher in abundance below than in the SMTZ (Fig. 3). We recovered one ANME-2a MAG (US2_17) from the newly categorized genus *Candidatus* Methanoborealis (Wallenius et al., 2026) that, in line with 16S rRNA amplicon reads, was more abundant below the SMTZ (0.8%) when compared to the SMTZ (0.2%).

We screened the MAGs with > 0.2 % abundance at either depth for marker genes in S-cycling, C_1_-cycling and potential for extracellular electron transport (EET) or metal reduction/oxidation (Fig. 3). S-cycling potential was widespread across the MAGs, with dissimilatory sulfate reduction genes *dsrA* and/or *aprA* found in multiple MAGs that were more abundant below the SMTZ, such as the two Desulfobacterales MAGs. However, the sulfur-oxidizing Sulfomarinibacterales MAG (US2_11) also encoded these genes and was more abundant in the SMTZ. In contrast, the sulfide:quinone oxidoreductase *sqr* was found in MAGs that were more abundant in the SMTZ and potentially used for sulfide detoxification. C1- cycle genes were scarce, indicating that the sediments harbor mainly heterotrophic clades. Only two MAGs with the marker gene for (reverse) methanogenesis, *mcrA,* were present; ANME-2a (*Ca.* Methanoborealis) and methanogenic Methanoregula (US2-13). Putative genes involved in EET and metal reduction/oxidation were found in multiple MAGs, which were often more abundant in the SMTZ (Fig. 3). Woeseiaceae MAG (US2_43) encoded multiple cytochrome c or membrane proteins involved in electron transfer across the membranes, i.e. RnfC from a complex that translocates proteins across the membrane and MtrA, MtrB and MtrC from the decaheme-cytochrome c that transfers electrons to the outer membrane. OmcS, OmcF and OmcZ, outer membrane cytochromes linked to metal reduction via EET, were present in a few genomes such as Desulfobacterota MAGs. Desulfobacterales_UBA11574 that encoded OmcS, and the most abundant Deferrimicrobium that encoded OmcF. The ANME-2a genome also showed potential for EET and metal reduction as RnfC and OmcZ homologue were detected in their genome.

### 3.4 Metatranscriptomic analysis shows active ANME-2a in the SMTZ

To identify the metabolically active members of the community that contribute to biogeochemical cycling, metatranscriptomes were sequenced from three replicates for each of the two depths. The analysis of mapped coding DNA sequence (CDS) reads revealed that the most active taxa were notably different from the most abundant taxa identified in the metagenomic DNA data (Fig. 3 and 4) as has been observed in other studies (e.g. Zhang et al., 2020). The proportion of unmapped reads was even greater in the metatranscriptomic dataset than in the metagenomic dataset, as ∼71 – 75 % of all transcripts did not map to the medium-quality MAGs analyzed. In the SMTZ, the most transcripts mapped to Aminicenantia_bin2 (MAG US2_47), which covered 12-22% of all CDS reads, and Xanthomonadales_bin3 (US2_56) with 7-8% relative abundance. Three putative SRBs, Desulfobacterales_JAFDCJ01 (US2_46), Desulfobacterales_UBA11574 (US2_27) and Desulfurivibrionaceae (US2_55) covered ∼6 % of the reads in the SMTZ, although they were either not in the top taxa at all or more abundant below the SMTZ in the metagenomic reads (Fig. S6). The *Ca.* Methanoborealis (ANME-2a) MAG had higher expression levels in the SMTZ than below the SMTZ with > 6 % and < 1% relative abundance, respectively (Fig. S7).

**Fig. 4.**
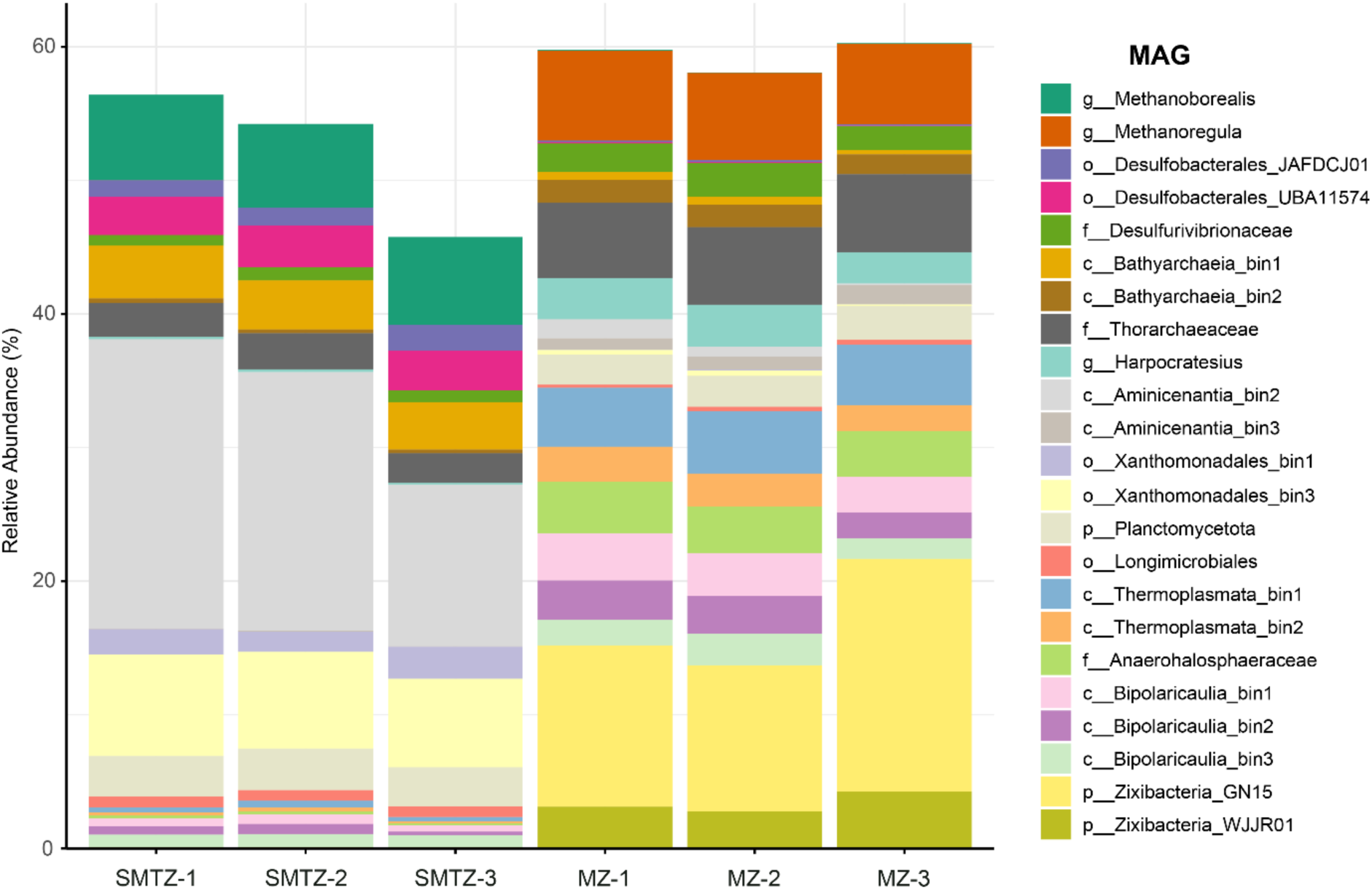
The relative abundance of expressed coding DNA sequence (CDS) reads of top MAGs at both depths (SMTZ: 16-20 cm; below the SMTZ in methanogenesis zone (MZ): 28-32 cm). Three replicates from each depth were sequenced for total RNA. The relative abundance here represents the percentage of total counts of coding genes per MAG detected in the transcriptome of each depth.

The active community below the SMTZ differed greatly from that in the SMTZ (Fig. 4). In general, archaeal MAGs were much more abundant in the deeper layer, such as Bathyarchaeia and Thermoplasmata bins. However, almost a third of the transcripts (∼20%) were covered by two Zixibacteria MAGs which were barely present in the SMTZ (0.001-0.2%).

### 3.5 Increased methane oxidation upon addition of electron acceptors

The potential of methane oxidation coupled to oxygen, sulfate, Mn and Fe oxide and graphene oxide, as an analogue for natural organic matter, was tested in sediment both from within the SMTZ (16 - 20 cm) and below the SMTZ (28 - 32 cm; Fig. 5). All incubations supplied with extra electron acceptor showed an increase in ^13^C-CO_2_/^12^C-CO_2_ ratio relative to the control where no electron acceptor was added, indicating active methane oxidation. Generally, the signal for methane oxidation was largest in the sediment from below the SMTZ (Fig. 5 and fig S8). Oxygen was the most effective electron acceptor added, inducing a methane oxidation rate of around 216 µmol cm^-3^ yr^-1^ at both depths (Fig. 5 and Fig. S8). After oxygen, the highest methane oxidation rates were obtained with sulfate as electron acceptor with 28.6 ± 0.4 and 8.0 ± 0.8 µmol cm^-3^ yr^-1^ for within and below the SMTZ, respectively. The methane oxidation rates in the Mn-, Fe- and NOM-AOM incubations were within a similar range, varying from 1.2 to 3.3 µmol cm^-3^ yr^-1^ for sediments in the SMTZ and from 9.4 to 15.5 µmol cm^-3^ yr^-1^ for those below the SMTZ.

**Fig. 5.**
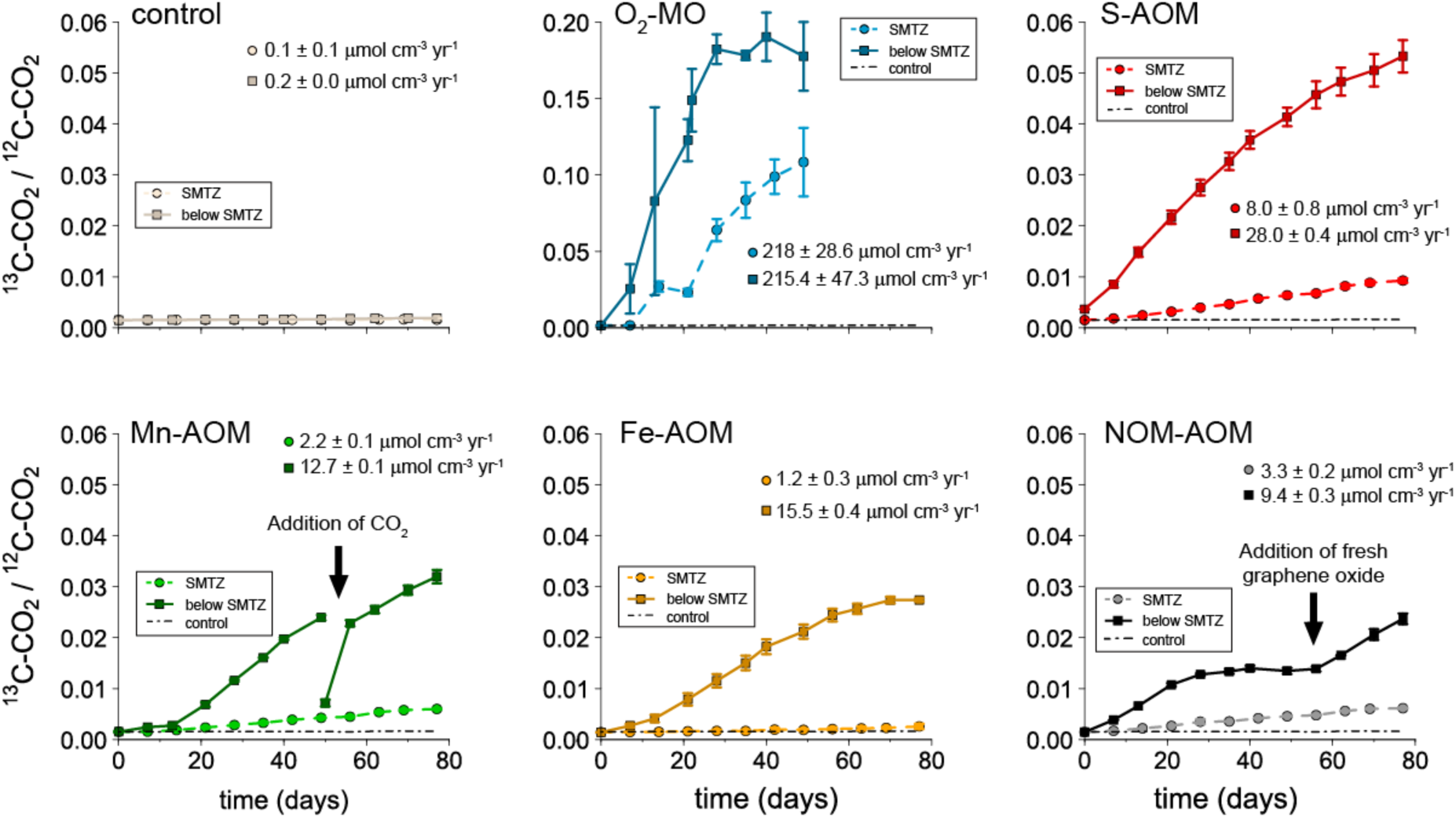
^13^C-CO_2_/^12^C-CO_2_ ratios in the headspace of the incubation experiments for sediment from within the SMTZ (16-20 cm; circles) and from below the SMTZ (28-32 cm; squares). The CH_4_ oxidation rates are based on the increase in ^45^C-CO_2_ in the headspace and liquid of the incubations (Fig. S8). Note the different y-axis at the plot showing the O_2_-MO incubations.

Accumulation of sulfate occurred in the O_2_-MO (max. 2.7 mmol L^-1^ within the SMTZ and 1.1 mmol L^-1^ below the SMTZ), Mn-AOM (max. 1 mmol L^-1^ within the SMTZ and 0.3 mmol L^-1^ below the SMTZ) and NOM-AOM (max. 0.3 mmol L^-1^ within the SMTZ and 0.5 mmol L^-1^ below the SMTZ) incubations (Fig. S9). In the controls and Fe-AOM incubations, no sulfate accumulated. Accumulation of sulfide was only observed in the S-AOM incubation from below the SMTZ (Fig. S9).

In the incubations where Mn oxide was added, dissolved Mn concentrations increased up to around 2 mmol L^-1^ within the first 30 days and subsequently stabilized for sediment from both depths (Fig. S9). Determination of the redox state of the dissolved Mn in incubations where Mn oxides were added showed that the accumulated dissolved Mn was almost exclusively Mn(II), indicating that reduction of Mn(IV) to Mn(II) occurs in one step and likely does not form dissolved Mn(III) as a reactive intermediate (Fig. S9). In the incubations with O_2_ as electron acceptor, accumulation of dissolved Mn up to a concentration of 0.8 mmol L^-1^ was also observed (Fig. S9). The other incubations had relatively stable dissolved Mn concentrations varying between 0.2 and 0.4 mmol L^-1^.

In the incubations with Fe oxides, dissolved Fe increased from 0.5 to 0.9 mmol L^-1^ in the SMTZ and from 0.2 to 0.5 mmol L^-1^ in the incubation with sediment from below the SMTZ (Fig. S9). When O_2_ was added as an electron acceptor, dissolved Fe accumulated to values of up to 1.1 mmol L^-1^ and 0.6 mmol L^-1^ in the incubations with sediment from within and below the SMTZ, respectively. In the other incubations, dissolved Fe was relatively stable or absent, apart from the incubations with graphene oxide, where the dissolved Fe concentrations fluctuated around 0.2 mmol L^-1^.

### 3.6 ANME archaea drive methane oxidation in the AOM incubations

ANME-2ab were, based on 16S rRNA analysis, the most abundant archaeal taxa in the incubations with the deeper sediment layer, covering ∼55 % of the total archaeal reads (Fig. S10). Since all ANME in the metagenomic analysis of the source sediment belong to ANME-2a (Table S4), we assume that all ANME in the incubation are ANME-2a. In the anoxic incubations, ANME reads increased significantly in the electron acceptor amended samples, but not in the control samples, underlining their role in AOM (Fig 6a). Upon addition of Mn and Fe oxides, ANME-2a increased by more than 20% points, accounting for up to 75 % of the relative archaeal abundance. *Methanosarcina* was the second most abundant archaea in all samples and increased the most in the control and graphene oxide treatments (3-fold and 2-fold, respectively).

**Fig. 6.**
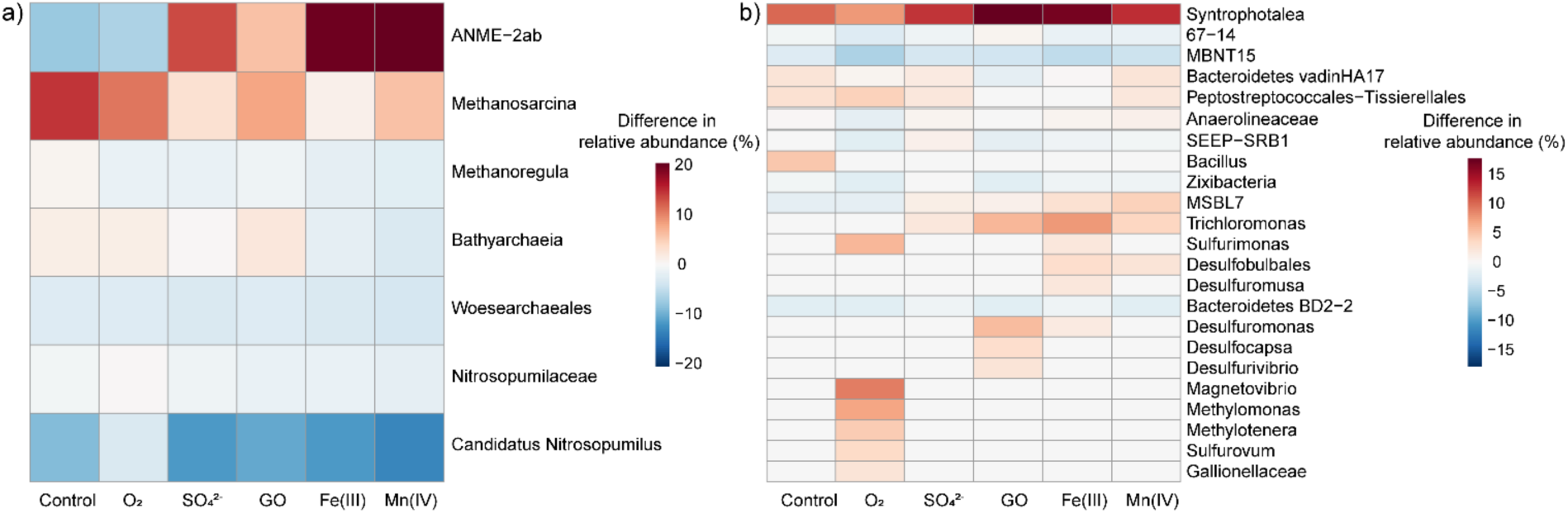
Changes in the relative abundance of microbial taxa in the sediment from below the SMTZ (28-32 cm) after the incubation with different electron acceptors compared to that at the start for (a) archaea and (b) bacteria. GO is used as an abbreviation for graphene oxide.

The 16S rRNA gene amplicon analysis revealed distinct differences between the microbial community of incubations with oxygen versus other electron acceptors (Fig. 6). With oxygen, the largest changes in the bacterial community were seen in *Methylomonas* and *Methylotenera* which increased in relative abundance by 5-10%, indicating that dormant buried cells were activated upon oxygen amendment (Fig. 6b). In addition, potential sulfur oxidizing taxa *Sulfurimonas*, *Sulfurovum* and *Magnetovibrio* increased in these samples. Interestingly, the changes in the bacterial community composition were highly substrate-dependent, although *Syntrophotalea* reads, of phylum Desulfobacterota, increased in all samples from <0.01% by 7-18%, the biggest increase was observed in the Fe-AOM and NOM-AOM treatments (Fig. 6b and Fig. S11). Also, other Desulfobacterota taxa increased, with Trichloromas and MSBL7 increasing in all but the control samples, whereas changes in other genera were more substrate-specific. Curiously, SEEP-SRB1, a putative SRB partner for ANME, increased only with sulfate, whereas other putative AOM syntrophs, uncharacterized Desulfobulbales, increased in Mn-AOM and Fe-AOM samples. The changes in heterotrophic bacteria involved in organic matter breakdown via fermentation, such as Bacteroidetes, were also substrate-specific (Fig. 6b).

## 4. Discussion

### 4.1 Two Distinct Zones of AOM

Our integrated analysis of sediments from the Bothnian Sea site reveals two vertically distinct zones of anaerobic oxidation of methane (AOM) which are both characterized by the presence of a metabolically versatile community of ANME-2a archaea but differ in their porewater and sediment chemistry. The first is the canonical sulfate-methane transition zone (SMTZ), where AOM is primarily coupled to sulfate reduction. Here, multiple indicators point towards AOM *in-situ*. The second is a deeper, previously unrecognized zone below the SMTZ, where ANME-2a persist and have the potential to couple AOM to a wide range of electron acceptors, although their *in-situ* activity appears to be limited. This finding is supported by a combination of geochemical profiling, DNA and RNA sequencing, and sediment incubation experiments, as discussed in detail below.

Active AOM in the SMTZ, which is located between depths of 12 and 22 cm (Fig. 1), is evident from the strong isotopic enrichment of ^13^C and D in the residual methane and depletion of ^13^C in DIC (Whiticar, 1999). This is corroborated by a peak in sulfate reduction and change in the gradient in porewater sulfate at the lower boundary of this zone, which is a typical feature in sediments where S-AOM occurs (Iversen & Jorgensen, 1985). Molecular analysis confirms these observations, with a high relative abundance of ANME-2a archaea (a maximum of 31% of archaeal sequences) co-occurring with putative partner sulfate-reducing bacteria from the *Desulfosarcinaceae* family (Schreiber et al., 2010; Murali et al., 2023; Fig. 2). Furthermore, metatranscriptomic data reveal that the *Ca.* Methanoborealis gene transcripts are highly expressed within the SMTZ, accounting for over 6% of mapped transcripts. Besides sulfate, the SMTZ at our site also contains Mn and Fe oxides and abundant organic matter (Fig. 1), which may all act as potential electron acceptors in AOM (Fig. 5). The addition of sulfate, Mn oxides, Fe oxides, and graphene oxide to sediment incubations all stimulated AOM at rates comparable to previous incubation studies with sulfate (range of 2 - 52 µmol CH_4_ cm^-3^_sed_ yr^-1^; Beal et al., 2009; Segarra et al., 2013; Aromokeye et al., 2020), Fe oxides (range of 0.035 - 6 µmol CH_4_ cm^-3^_sed_ yr^-1^; Beal et al., 2009; Segarra et al., 2013; Egger et al., 2015; Aromokeye et al., 2020; Xu et al., 2021) and Mn oxides (range of 0.20 - 14 µmol CH_4_ cm^-3^_sed_ yr^-1^; Beal et al., 2009; Segarra et al., 2013; Xu et al., 2021). However, the accumulation of sulfate in the incubations amended with Mn and graphene oxide makes it difficult to distinguish direct metal-AOM from AOM coupled to a cryptic sulfur cycle in these treatments (e.g. Holmkvist et al., 2011; Su et al., 2020). Taken together, these results provide strong indications that ANME-2a are the primary microorganisms actively mediating AOM in the SMTZ, with S-AOM as the dominant pathway.

More surprisingly, we identified a second, deeper zone of potential AOM below the SMTZ. Here, the relative abundance of ANME-2a is even higher, peaking at 61% of the archaeal community between 24 and 34 cm. This zone is depleted in sulfate but contains solid phase Mn and, in a few depth intervals, Fe oxides that have been buried below the current SMTZ. The methane isotope profile does not show a clear signal for oxidation in this deeper layer (Fig. 1), but an AOM signal could be masked by concurrent methanogenesis (Seifert et al., 2006), which, based on the relative abundance of methanogens, occurs below a depth of 28 cm (Fig. 2).

Direct observations for the metabolic potential of this deeper community are derived from our sediment incubation experiments. The addition of sulfate, Mn oxides, Fe oxides and graphene oxides all stimulated AOM, with consistently higher rates compared to the sediment within the SMTZ (Fig. 5), possibly linked to a higher initial ANME biomass (Fig. 2 and Fig. S6). In every anaerobic incubation for the sediment interval from 28-32 cm, the relative abundance of ANME-2a increased significantly, particularly in the presence of metal oxides, where they finally constituted up to 75% of the archaeal community (Fig. 6). The sediment incubations indicate that the ANME-2a are capable of oxidizing methane using sulfate and the alternative electron acceptors tested, although again a cryptic sulfur cycle cannot always be excluded. Notably, the lack of sulfate accumulation in the Fe-amended incubations points towards a direct coupling between AOM and Fe reduction. Metagenomic analyses on the genome of ANME-2a (*Ca*. Methanoborealis) show a high potential for metal-AOM (Wallenius et al., 2026). Direct metal-AOM by ANME-2a is also supported by our genomic data, which show that the ANME-2a MAG recovered from this site encodes a homologue for OmcZ (Fig. 3), a multiheme cytochrome known to be involved in extracellular electron transfer to metals in bacteria (Gu et al., 2023), although its function in ANME is not confirmed. Notably, the ANME-2a in this deeper community were detectable through metatranscriptomics (Fig. S7), although their activity was lower compared to the community within the SMTZ. This strongly suggests that this deep ANME-2a population is not dormant.

### 4.2 Environmental relevance

The existence of a deep, active ANME-2a community may be explained by the recent environmental history of the site. Geochemical evidence suggests that the lower boundary of the SMTZ has become shallower over the past decades. This is based on a previously observed depth of approximately 28 cm for the lower boundary of the SMTZ at this site (Slomp et al., 2013) and on the mismatch between the current position of the SMTZ and the FeS and total S peak in the sediment (Fig. 1). In a setting where sulfide is largely restricted to the SMTZ or, as is the case here, is absent from the porewater, such solid phase sulfur peaks are expected within the SMTZ and not below (Riedinger et al., 2014; Egger et al., 2015). The upward shift of the SMTZ at our study site has left a large population of ANME-2a stranded in a now sulfate-depleted environment. Our data suggest that this community has remained active using one or more solid phase electron acceptors, possibly partly through a cryptic sulfur cycle, thereby establishing a second, deeper microbial filter for methane. Such a low rate of AOM fueled by other electron acceptors than sulfate could also explain the presence of ANME-2a below the SMTZ at sites where upward shifts in SMTZ depth have not occurred recently (Aromokeye et al., 2020; Deng et al., 2020; Dalcin Martins et al., 2024). In conclusion, a deep microbial methane filter, driven by ANME-2a below the SMTZ, could be a general feature in many coastal sediments.

Our findings have significant implications for our understanding of coastal methane cycling. Eutrophication is expected to continue to increase globally (Breitburg et al., 2018). Variations in river runoff due to climate change may lead to a fluctuating supply of organic matter and metal oxides to coastal systems, especially at high latitudes (Canuel et al., 2012; Lenstra et al., 2018). Combined, this may lead to a large variability in the depth of the SMTZ and the availability of metal oxides, and potentially, humic substances, to methanotrophs. The versatility of ANME-2a in using these electron acceptors is expected to contribute to a more resilient methane filter in coastal brackish sediments in two ways. First, a substantial amount of methane could be removed by ANME below the SMTZ via AOM pathways other than S-AOM, given the relatively large overlap of methane with electron acceptors like Fe and Mn oxides at many coastal locations (Egger et al., 2015; Aromokeye et al., 2020; Lenstra et al., 2023; Xiao et al., 2023). Second, a downward shift of the SMTZ, as predicted by reactive transport models upon a decrease in organic matter supply (Rooze et al., 2016), could directly enable S-AOM by the ANME-2a community deeper in the sediment. Methanotrophic archaea have notoriously low growth rates and S-AOM is thus often limited by ANME biomass (Dale et al., 2006; Lenstra et al., 2023). Hence, a rapid start of S-AOM by ANME-2a upon a downward shift of the SMTZ, could increase the efficiency of the methane filter.

The persistence and versatility of the ANME-2a community in using a range of electron acceptors in AOM should be incorporated in computer models designed to predict methane dynamics in coastal sediments (Lenstra et al., 2023; Wallheimer et al., 2025). Future research should also focus on quantifying the *in-situ* rates of Mn-, Fe- and NOM-driven AOM in brackish coastal sediments and in determining the global distribution of such AOM systems to refine estimates and models of the marine methane budget in a changing world (Wallenius et al., 2021; Saunois et al., 2025).

## Supporting information

Supplemental data Klomp, Wallenius et al

## Data availability

The dataset for this study is available through Zenodo at https://doi.org/10.5281/zenodo.16920164. Sequencing data is available at the National Center for Biotechnology Information (NCBI) website under the accession number PRJNA1306556.

## Acknowledgements

We thank the captain and crew, Anna Palmbo Bergman and Nina Dagberg for assistance during sampling aboard the KBV 181. We thank Anita van Leeuwen, Arnold van Dijk, Carina van Veen, Coen Mulder, John Visser, Sebastian Krosse and Thom Claessen for analytical assistance in the lab. This research was funded by ERC Synergy grant MARIX 854088 and NESSC NWO-OCW 024002001.

