## Supplemental data Klomp, Wallenius et al for "ANME-2a drive methane oxidation in brackish coastal sediments via multiple pathways"

### Supplementary figures

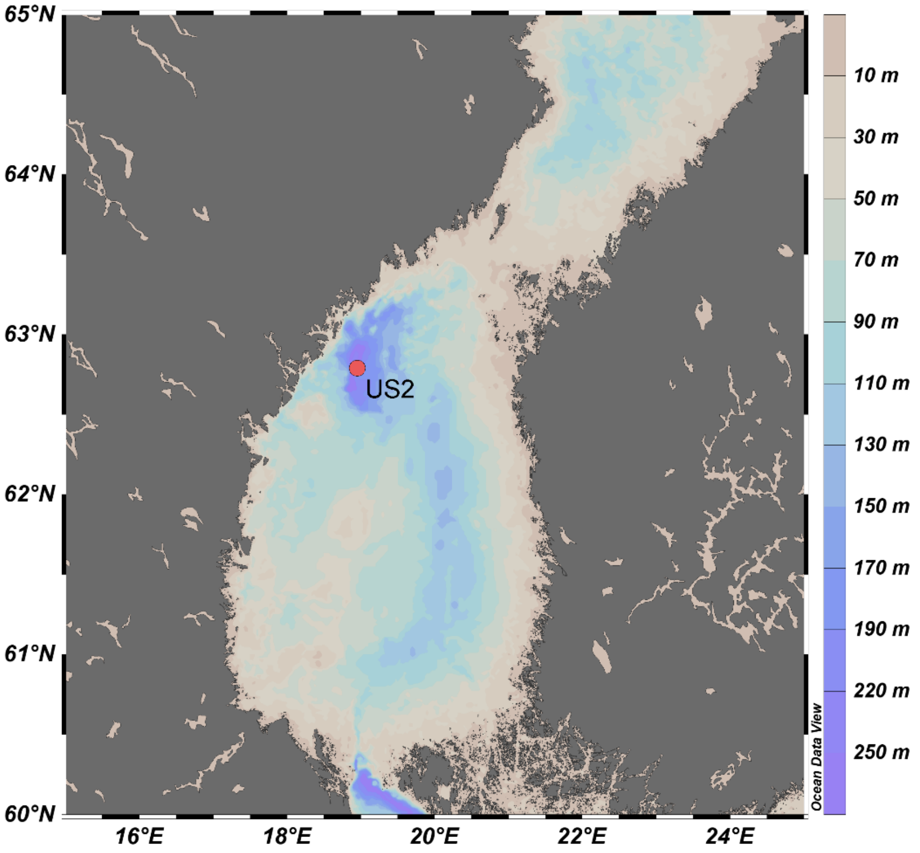


**Fig. S1.** Map of the Bothnian Sea, including the location of site US2. Figure obtained with Ocean Data View (Schlitzer, 2023).

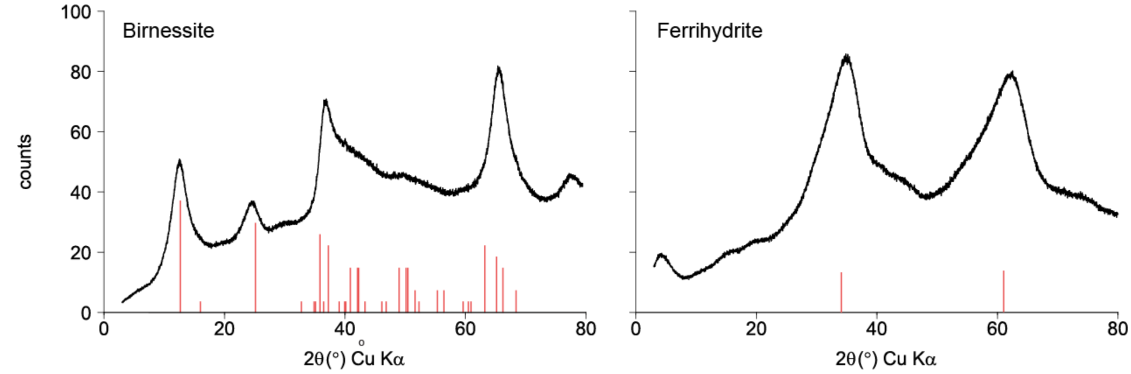


**Fig. S2** X-ray diffraction spectra of birnessite and ferrihydrite. The red bars indicate the theoretical positions of the 2θ peaks for birnessite (Lenstra et al., 2021) and ferrihydrite (Das et al., 2011).

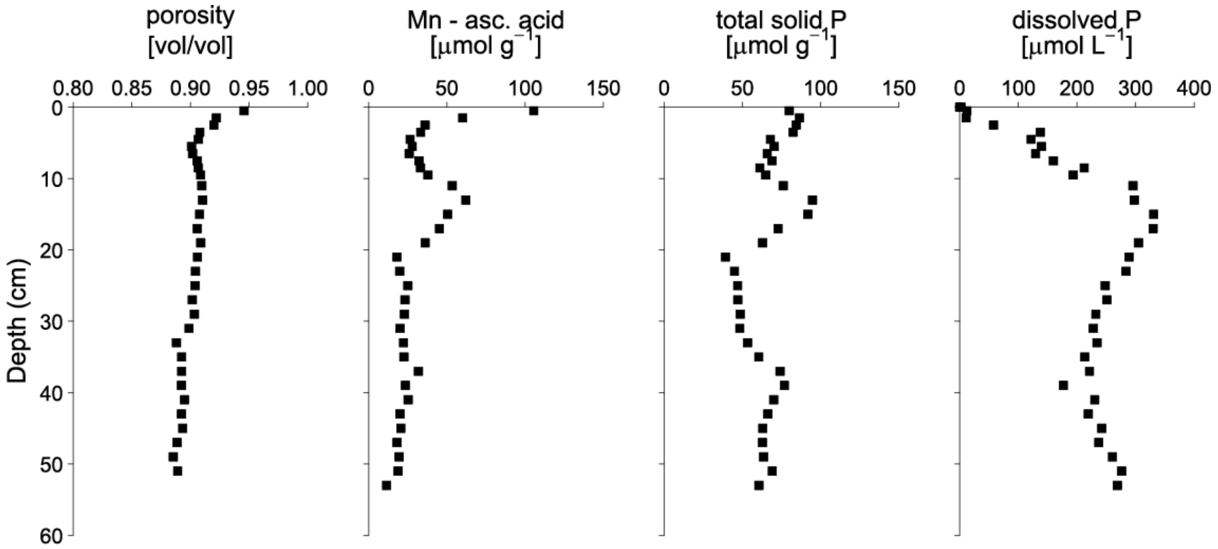


**Fig. S3.** Sediment profiles of porosity, Mn extracted in ascorbic acid, total solid P and dissolved P. The maximum in Mn extracted in ascorbic acid between depths of 10 and 20 cm corresponds to a maximum in total solid P, which suggests that the peak represents a Mn phosphate phase. This zone also corresponds with the maximum dissolved P concentrations in the pore water.

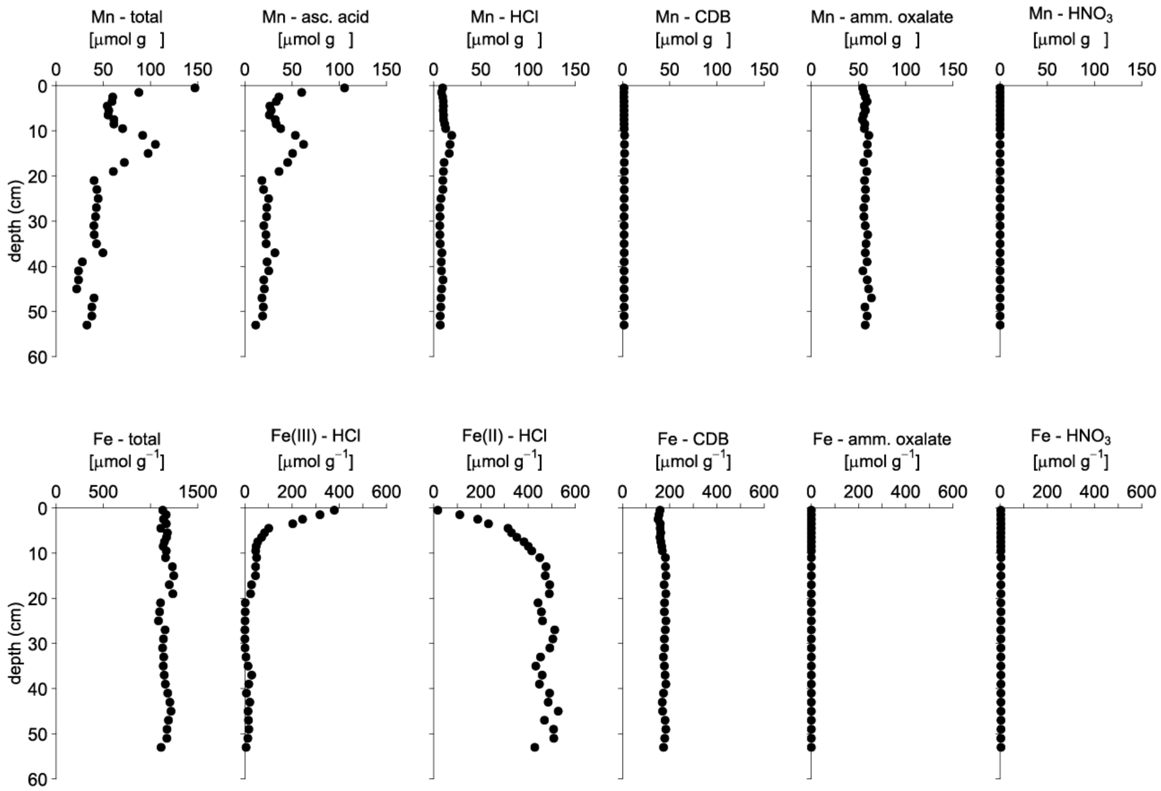


**Fig. S4.** Sediment profiles of total Mn and Fe and from all steps in the sequential extractions. Ascorbic acid (asc. acid) extracts poorly ordered Mn oxides, HCl extracts Mn carbonates and easily reducible Fe(III) and Fe(II) minerals, sodium dithionite (CDB) extracts crystalline Fe and Mn oxides and Mn bound to clays, ammonium oxalate (amm. oxalate) extracts recalcitrant Fe oxides, crystalline Mn oxides and Mn bound to clays and $HNO_3$ extracts pyrite, including any associated Mn. Note the different range for the x-axis of Fe total.

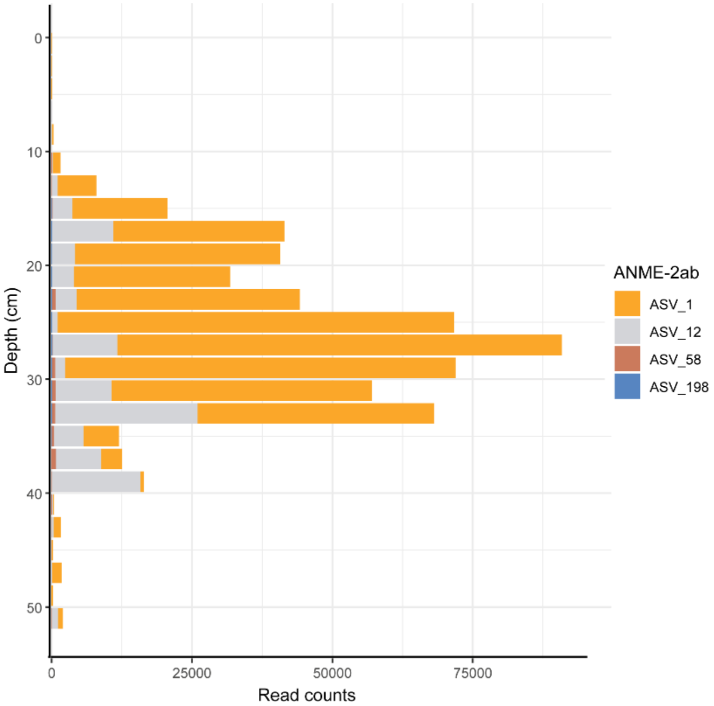


**Fig. S5.** The total read counts of different ASVs assigned taxonomically to ANME-2ab across the sediment depths.

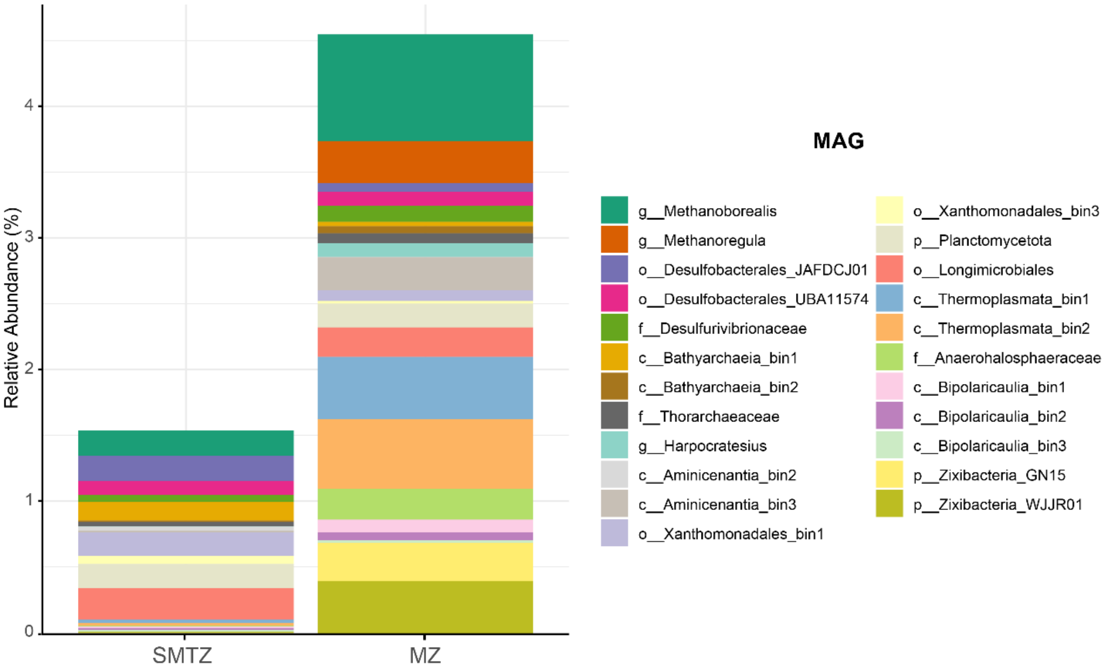


**Fig. S6.** The MAGs with the highest coverage in metatranscriptome data (Fig. 4) in the SMTZ and below the SMTZ in the methanogenic zone (MZ), portrayed as relative abundance of reads mapped to the metagenome of each depth.

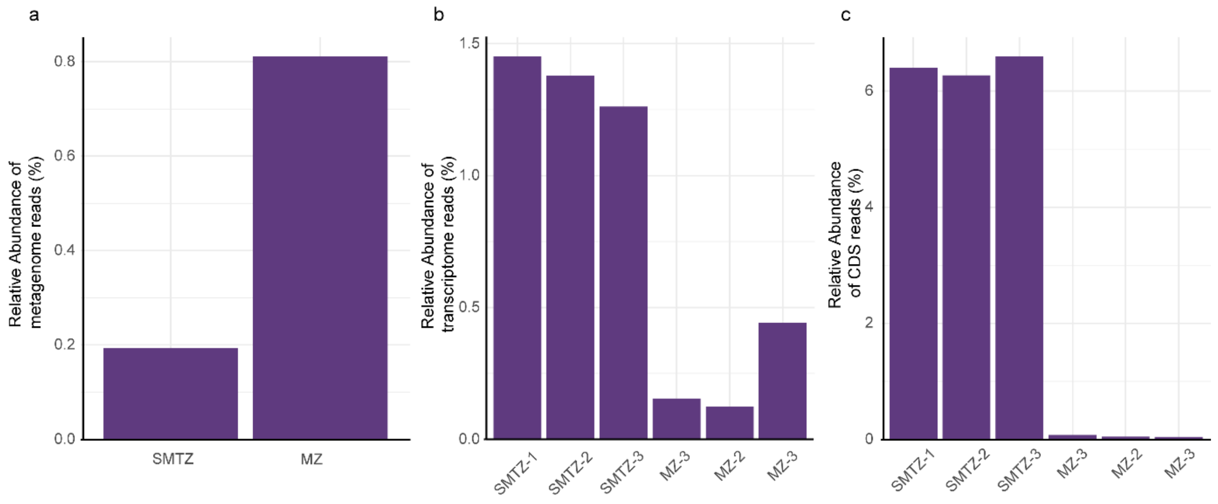


**Fig. S7.** The abundance of ANME-2a MAG when mapped to a) the whole metagenome, b) the whole transcriptome; and c) the relative abundance of ANME-2a MAG CDS reads from all CDS reads mapped to analyzed MAGs in both the SMTZ and below the SMTZ in the methanogenic zone (MZ), portrayed as relative abundance of reads mapped to the metagenome of each depth.

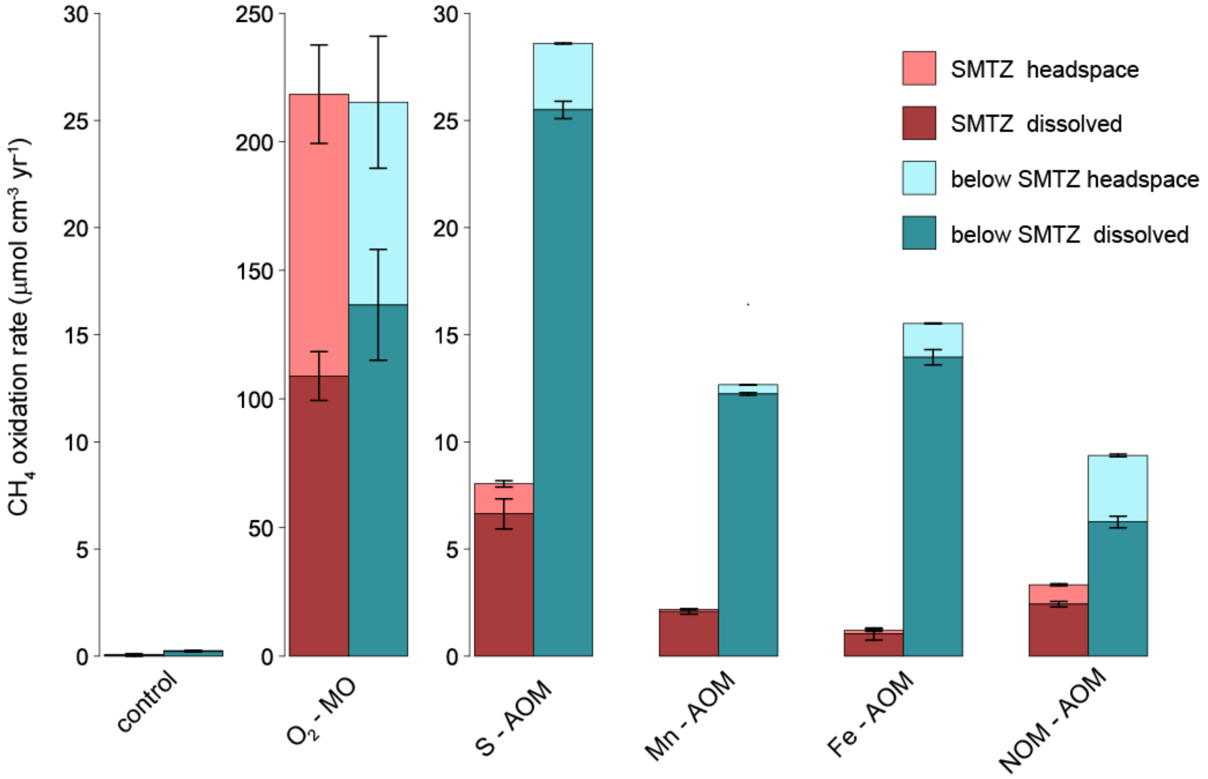


**Fig. S8.** Maximum methane oxidation rates in the different incubation experiments of sediment from within and below the SMTZ, calculated based on the increase in $^{13}C$-$CO_2$ and $^{12}C$-$CO_2$ in the headspace (lighter color) and the dissolved phase (darker color).

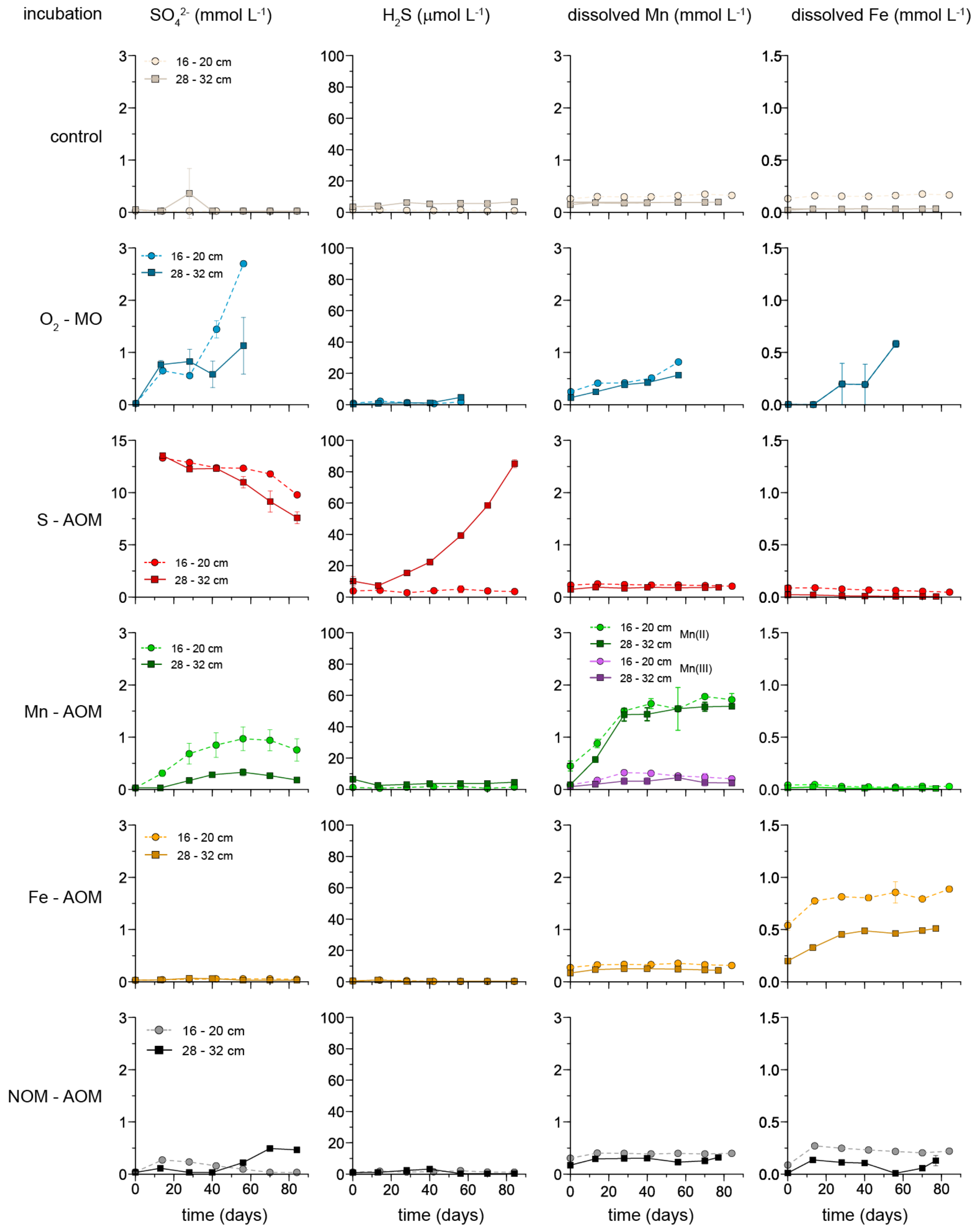


**Fig. S9.** Concentrations of sulfate, sulfide, dissolved Mn and dissolved Fe in the incubations. Note that, in the Mn-AOM incubation, the redox speciation of dissolved Mn was determined, providing insight in concentrations of Mn(II) and Mn(III).

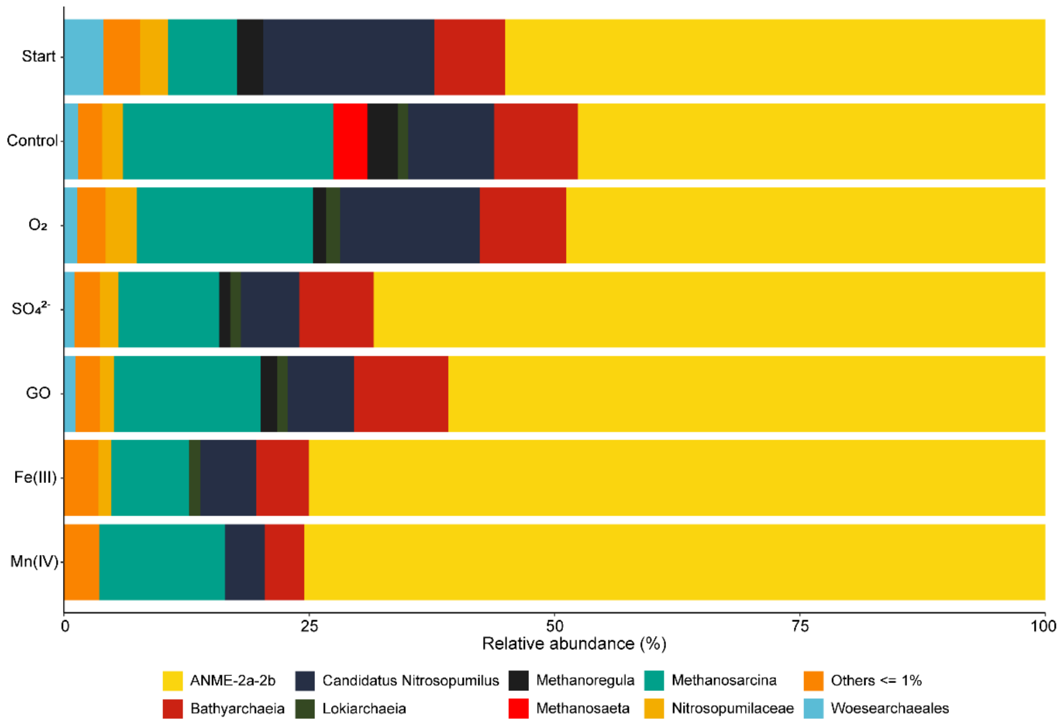


**Fig. S10.** The relative abundance of archaeal taxa with >1% abundance in all AOM incubations and in the sediment at the start.

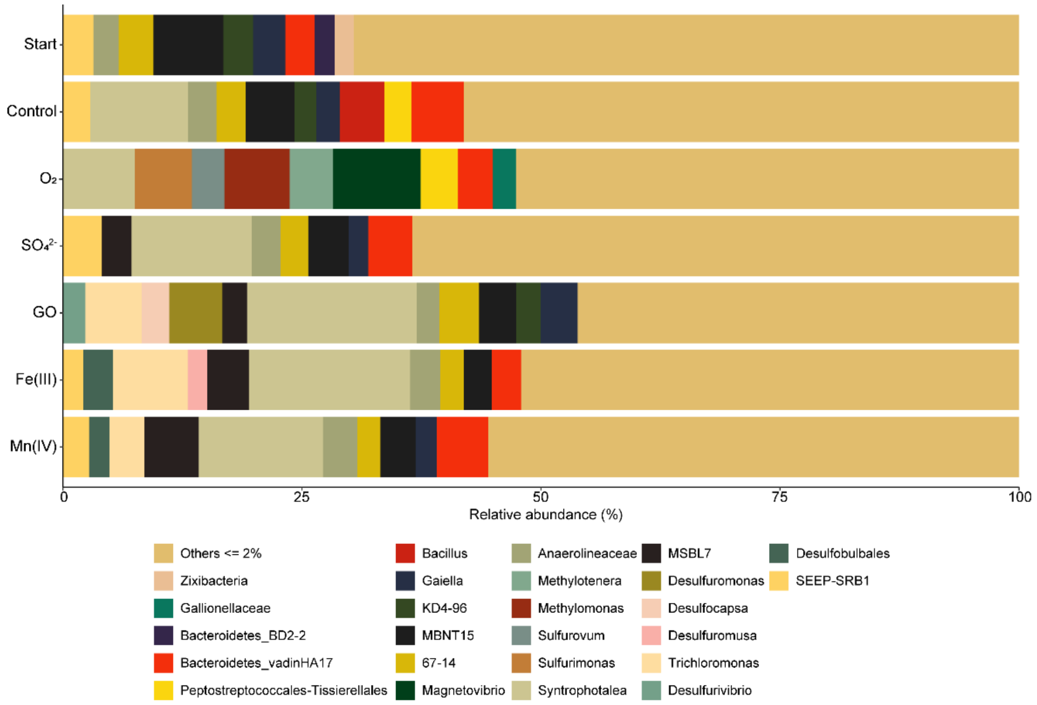


**Fig. S11.** The relative abundance of bacterial taxa with >1% abundance in all AOM incubations and in the sediment at the start.

**Supplementary tables**

**Table S1.** Scheme for the sediment Mn extraction (Lenstra et al., 2021)

| Step | Extraction solution | Time of extraction (h) | Mineral phase targeted |
|---|---|---|---|
| 1 | 0.17 M sodium citrate, 0.6 M sodium bicarbonate and 0.057 M ascorbic acid (pH 7.5) | 24 | Mn oxide and Mn phosphates |
| 2 | 1 M HCl | 4 | Mn carbonate and Mn sulfides |
| 3 | 50 g $L^{-1}$ sodium dithionite solution buffered with 0.35 M acetic acid/0.2 M sodium citrate to pH 48 | 4 | Non-reactive Mn bound to clays |
| 4 | 0.2 M ammonium oxalate / 0.17 M oxalic acid (pH 3.2) | 6 | Non-reactive Mn bound to clays |
| 5 | 65% $HNO_3$ | 2 | Mn bound to and incorporated in pyrite |

**Table S2.** Scheme for the sediment Fe extraction (Kraal et al. (2017)

| Step | Extraction solution | Time of extraction (h) | Mineral phase targeted |
|---|---|---|---|
| 1 | 1 M HCl | 4 | Poorly ordered or pH-sensitive Fe(III) and Fe(II) minerals (e.g. ferrihydrite, siderite or Fe monosulfide) |
| 2 | 50 g $L^{-1}$ sodium dithionite solution buffered with 0.35 M acetic acid/0.2 M sodium citrate to pH 48 | 4 | Crystalline Fe oxides (goethite, hematite) |
| 3 | 0.2 M ammonium oxalate / 0.17 M oxalic acid (pH 3.2) | 6 | Recalcitrant Fe oxides (magnetite) |
| 4 | 65% $HNO_3$ | 2 | Pyrite |

**Table S3.** Composition of the artificial sulfate-free seawater (pH 7.5).

| Compound | Concentration |
|---|---|
| NaCl | 5.2 g $L^{-1}$ |
| $MgCl_2$ * 6 $H_2O$ | 1 g $L^{-1}$ |
| $CaCl_2$ * 2 $H_2O$ | 0.28 g $L^{-1}$ |
| KCl | 0.1 g $L^{-1}$ |
| HEPES | 20 mmol $L^{-1}$ |

**Table S4.** Taxonomy and relative abundance of the different ANME MAG's obtained in the metagenomic analysis of the sediment samples, based on SingleM analysis.

| Taxonomy | Incubation within SMTZ (16-20 cm) | Incubation below SMTZ (28-32 cm) |
|---|---|---|
| Root; d__Archaea; p__Halobacteriota; c__Methanosarcinia; o__Methanosarcinales; f__Methanocomedenaceae; g__Kmv04 | 0.37 | 1.71 |
| Root; d__Archaea; p__Halobacteriota; c__Methanosarcinia; o__Methanosarcinales; f__Methanocomedenaceae; g__QBUR01 | 0 | 0.01 |
| Root; d__Archaea; p__Halobacteriota; c__Methanosarcinia; o__Methanosarcinales; f__Methanocomedenaceae; g__Methanocomedens | 0 | 0 |
